# ML-DSP: Machine Learning with Digital Signal Processing for ultrafast, accurate, and scalable genome classification at all taxonomic levels

**DOI:** 10.1101/394932

**Authors:** Gurjit S. Randhawa, Kathleen A. Hill, Lila Kari

## Abstract

**Background:** Although methods and software tools abound for the comparison, analysis, identification, and taxonomic classification of the enormous amount of genomic sequences that are continuously being produced, taxonomic classification remains challenging. The difficulty lies within both the magnitude of the dataset and the intrinsic problems associated with classification. The need exists for an approach and software tool that addresses the limitations of existing alignment-based methods, as well as the challenges of recently proposed alignment-free methods.

**Results:** We combine supervised **M**achine **L**earning with **D**igital **S**ignal **P**rocessing to design **ML-DSP**, an alignment-free software tool for ultrafast, accurate, and scalable genome classification at all taxonomic levels.

We test ML-DSP by classifying 7,396 full mitochondrial genomes from the kingdom to genus levels, with 98% classification accuracy. Compared with the alignment-based classification tool MEGA7 (with sequences aligned with either MUSCLE, or CLUSTALW), ML-DSP has similar accuracy scores while being significantly faster on two small benchmark datasets (2,250 to 67,600 times faster for 41 mammalian mitochondrial genomes). ML-DSP also successfully scales to accurately classify a large dataset of 4,322 complete vertebrate mtDNA genomes, a task which MEGA7 with MUSCLE or CLUSTALW did not complete after several hours, and had to be terminated. ML-DSP also outperforms the alignment-free tool FFP (Feature Frequency Profiles) in terms of both accuracy and time, being three times faster for the vertebrate mtDNA genomes dataset.

**Conclusions:** We provide empirical evidence that ML-DSP distinguishes complete genome sequences at all taxonomic levels. Ultrafast and accurate taxonomic classification of genomic sequences is predicted to be highly relevant in the classification of newly discovered organisms, in distinguishing genomic signatures, in identifying mechanistic determinants of genomic signatures, and in evaluating genome integrity.

## Introduction

Of the estimated existing 8.7 million (±1.3 million) species existing on Earth [1], only around 1.5 million distinct eukaryotes have been catalogued and classified so far [2], leaving 86% of existing species on Earth and 91% of marine species still unclassified. To address the grand challenge of all species identification and classification, a multitude of techniques have been proposed for genomic sequence analysis and comparison. These methods can be broadly classified into alignment-based and alignment-free. Alignment-based methods and software tools are numerous, and include, e.g., MEGA7 [3] with sequence alignment using MUSCLE [4], or CLUSTALW [5, 6]. Though alignment-based methods have been used with significant success for genome classification, they have limitations [7] such as the heavy time/memory computational cost for multiple alignment in multigenome scale sequence data, the need for continuous homologous sequences, and the dependence on a priori assumptions on, e.g., the gap penalty and threshold values for statistical parameters [8]. In addition, with next-generation sequencing (NGS) playing an increasingly important role, it may not always be possible to align many short reads coming from different parts of genomes [9]. To address situations where alignment-based methods fail or are insufficient, alignment-free methods have been proposed [10], including approaches based on Chaos Game Representation of DNA sequences [11, 12, 13], random walk [14], graph theory [15], iterated maps [16], information theory [17], category-position-frequency [18], spaced-words frequencies [19], Markov-model [20], thermal melting profiles [21], word analysis [22], among others. Software implementations of alignment-free methods also exist, among them COMET [23], CASTOR [24], SCUEAL [25], REGA [26], KAMERIS [27], and FFP (Feature Frequency Profile) [28].

While alignment-free methods address some of the issues faced by alignment-based methods, [7] identified the following challenges of alignment-free methods:

i. `Lack of software implementation:` Most of the existing alignment-free methods are still exploring technical foundations and lack software implementation, which is necessary for methods to be compared on common datasets.
ii. `Use of simulated sequences or very small real world datasets:` The majority of the existing alignment-free methods are tested using simulated sequences or very small real-world datasets. This makes it hard for experts to pick one tool over the others.
iii. `Memory overhead:` Scalability to multigenome data can cause memory overhead in word-based methods, specially when long *k*-mers are used. Information-theory methods address this issue using compression algorithms, but they may fail in identifying complex organization levels in the sequences.

In our effort to overcome these challenges, we propose ML-DSP (supervised **M**achine **L**earning based on **D**igital **S**ignal **P**rocessing), a general-purpose alignment-free method and software tool for genomic DNA sequence classification at all taxonomic levels.

### Numerical representations of DNA sequences

Digital signal processing can be employed in this context because genomic sequences can be numerically represented as discrete numerical sequences and hence treated as digital signals. Several numerical representations of DNA sequences, that use numbers assigned to individual nucleotides, have been proposed in the literature[29], e.g., based on a fixed mapping of each nucleotide to a number without biological significance, using mappings of nucleotides to numerical values deduced from their physio-chemical properties, or using numerical values deduced from the doublets or codons that the individual nucleotide was part of [29, 30]. In [31, 32] three physio-chemical based representations of DNA sequences (atomic, molecular mass, and Electron-Ion Interaction Potential, EIIP) were considered for genomic analysis, and the authors concluded that the choice of numerical representation did not have any effect on the results. The latest study comparing different numerical representation techniques [33] concluded that multi-dimensional representations (such as Chaos Game Representation) yielded better genomic comparison results than one-dimensional representations. However, in general there is no agreement on whether or not the choice of numerical representation for DNA sequences makes a difference on the genome comparison results, or what are the numerical representations that are best suited for analyzing genomic data. We address this issue by providing a comprehensive analysis and comparison of thirteen one-dimensional numerical representations for suitability in genome analysis.

### Digital Signal Processing

Following the choice of a suitable numerical representation for DNA sequences, digital signal processing (DSP) techniques can be applied to the resulting discrete numerical sequences, and the whole process has been termed genomic signal processing [30]. DSP techniques have previously been used for DNA sequence comparison, e.g., to distinguish coding regions from non-coding regions [34, 35, 36], to align the genomic signals for classification of biological sequences [37], for whole genome phylogenetic analysis [38], and to analyze other properties of genomic sequences [39]. In our approach, genomic sequences are represented as discrete numerical sequences, treated as digital signals, transformed via Discrete Fourier Transform into corresponding magnitude spectra, and compared via Pearson Correlation Coefficient to create a pairwise distance matrix.

### Supervised Machine Learning

Machine learning has been successfully used in small-scale genomic analysis studies [40, 41, 42]. In this paper we propose a novel combination of supervised machine learning with feature vectors consisting of the distance between the magnitude spectrum of a sequence’s digital signal and the magnitude spectra of all other sequences in the training set. The taxonomic labels of sequences are provided for training purposes. Six supervised machine learning classifiers (Linear Discriminant, Linear SVM, Quadratic SVM, Fine KNN, Subspace Discriminant, and Subspace KNN) are trained on this pairwise distance vectors, and then used to classify new sequences. Independently, classical multidimensional scaling generates a 3D visualization, called Molecular Distance Map (MoDMap) [43], of the interrelationships among all sequences.

For our computational experiments, we used a large dataset of 7, 396 complete mtDNA sequences, and six different classifiers, to compare one-dimensional numerical representations for DNA sequences used in the literature for classification purposes. For this dataset, we concluded that the Purine/Pyrimidine (PP), Real, and Just-A numerical representations were the top three performers. We analyzed the performance of ML-DSP in classifying the aforementioned genomic mtDNA sequences, from the highest level (domain into kingdoms) to lower level (family into genera) taxonomical ranks. The average classification accuracy of the ML-DSP was 98% when using the PP, Real, and Just-A numerical representations.

To evaluate our method, we compared its performance (accuracy and speed) on three datasets: two previously used small benchmark datasets [44], and a large real world dataset of 4, 322 complete vertebrate mtDNA sequences. We found that MLDSP had significantly better accuracy scores than the alignment-free method FFP on the two small benchmark datasets, while having similar accuracy but better running time on the large benchmark dataset. When compared to the state-of-the-art alignment-based method MEGA7, with alignment using MUSCLE or CLUSTALW, ML-DSP achieved similar accuracy but superior processing times (2,250 to 67,600 times faster) for the small benchmark dataset of 41 mammalian genomes. For the large dataset, ML-DSP took 28 seconds, while MEGA7(MUSCLE/CLUSTALW) could not complete the computation after 2 hours/6 hours, and had to be terminated.

## Results and Discussion

Following the design and implementation of the ML-DSP genomic sequence classification tool prototype, we investigated which type of length-normalization and which type of distance were most suitable for genome classification using this method. We then conducted a comprehensive analysis of the various numerical representations of DNA sequences used in the literature, and determined the top three performers. Having set these three parameters (length-normalization method, distance, and numerical representation), we tested ML-DSP’s ability to classify mtDNA genomes at taxonomic levels ranging from the domain level down to the genus level, and obtained average levels of classification accuracy of 98%. Finally, we compared ML-DSP with other alignment-based and alignment-free genome classification methods, and showed that ML-DSP achieved higher accuracy and significantly higher speeds.

### Analysis of distances and of length normalization approaches

To decide which distance measure and which length normalization method were most suitable for genome comparisons with ML-DSP, we used nine different subsets of full mtDNA sequences from our dataset. These subsets were selected to include most of the available complete mtDNA genomes (Vertebrates dataset of 4,322 mtDNA sequences), as well as subsets containing similar sequences, of similar length (Primates dataset of 148 mtDNA sequences), and subsets containing mtDNA genomes showing large differences in length (Plants dataset of 174 mtDNA sequences).

The classification accuracy scores obtained using the two considered distance measures (Euclidean and Pearson Correlation Coeffient, PCC) and two different length-normalization approaches (normalization to maximum length and normalization to median length) on several datasets are listed in Table 1. The classification accuracy scores are similar, which indicates that the choice of Euclidean or PCC, and the choice of one of the above length-normalization approaches do not result in a large difference in accuracy scores.

**Table 1.** Maximum classification accuracy scores when using Euclidean vs. Pearson’s correlation coefficient (PCC) as a distance measure.

| Data Set | No. of Seq. | Max Length (bp) | Min Length (bp) | Median Length (bp) | Maximum Accuracy |  |  |  |
| --- | --- | --- | --- | --- | --- | --- | --- | --- |
|  |  |  |  |  | Euclidean |  | PCC |  |
|  |  |  |  |  | Norm. to Max Length (a) | Norm. to Median Length (b) | Norm. to Max Length (c) | Norm. to Median Length (d) |
| Primates (Haplorrhini: 115, Strepsirrhini: 33) | 148 | 17531 | 15467 | 16554 | 100% | 100% | 100% | 100% |
| Protists (Alveolata: 34, Rhodophyta: 46, Stramenopiles: 79) | 159 | 77356 | 5882 | 35660 | 93.7% | 93.7% | 96.9% | 96.9% |
| Fungi (Basidiomycota: 30, Pezizomycotina: 104, Saccharomycotina: 92) | 226 | 235849 | 1364 | 39154 | 85.3% | 87.5% | 88.8% | 91.1% |
| Plants (Chlorophyta: 44, Streptophyta: 130) | 174 | 1999595 | 12998 | 128211 | 94.3% | 96.0% | 93.7% | 93.1% |
| Amphibians (Anura: 161, Caudata: 95, Gymnophiona: 34) | 290 | 28757 | 15757 | 17271 | 99.0% | 99.0% | 98.6% | 99.3% |
| Mammals (Xenarthrans: 30, Bats: 54, Carnivores: 135, Even-toed Ungulates: 242, Insectivores: 40, Marsupials: 34, Primates: 148, Rodents and Rabbits: 147) | 830 | 17734 | 15289 | 16537 | 99.3% | 99.4% | 98.9% | 99.0% |
| Insects (Coleoptera: 95, Dictyptera: 77, Diptera: 149, Hemiptera: 126, Hymenoptera: 47, Lepidoptera: 294, Orthoptera: 110) | 898 | 20731 | 10662 | 15529 | 96.3% | 97.1% | 96.1% | 97.0% |
| 3 classes (Amphibians: 290, Mammals: 874, Insects: 1006) | 2170 | 28757 | 8118 | 16361 | 99.9% | 99.8% | 99.8% | 99.8% |
| Vertebrates (Amphibians: 290, Birds: 553, Fish: 2313, Mammals: 874, Reptiles: 292) | 4322 | 28757 | 14935 | 16616 | 99.5% | 99.7% | 99.5% | 99.6% |
| **Table Average Accuracy** | — | — | — | — | 96.4% | 96.9% | 96.9% | 97.3% |

(a)(c) Genomes normalized to the maximum genome sequence length; (b)(d) Genomes normalized to the median genome sequence length

In the remainder of this paper we chose the Pearson Correlation Coefficient PCC because it is scale independent (unlike the Euclidean distance, which is, e.g., sensitive to the offset of the signal, whereby signals with the same shape but different starting points are regarded as dissimilar [45]), and the length-normalization to median length because it is economic in terms of memory usage.

### Analysis of various numerical representations of DNA sequences

We analyzed the effect on the classification accuracy of ML-DSP of the use of each of thirteen different one-dimensional numeric representations for DNA sequences, grouped as: Fixed mappings DNA numerical representations (rows 1, 2, 3, 6, 7, 10, 11, 12, 13, in Table 2), mappings based on some physio-chemical properties of nucleotides (rows 4, 5, in Table 2), and mappings based on the nearest-neighbour values (rows 8, 9, in Table 2).

**Table 2.** Numerical representations of DNA sequences.

| | Representation | Rules | Output for $S_1 = CGAT$ |
| --- | --- | --- | --- |
| 1 | Integer | $T = 0, C = 1, A = 2, G = 3$ | [1 3 2 0] |
| 2 | Integer (other variant) | $T = 1, C = 2, A = 3, G = 4$ | [2 4 3 1] |
| 3 | Real | $T = -1.5, C = 0.5, A = 1.5, G = -0.5$ | [0.5 -0.5 1.5 -1.5] |
| 4 | Atomic | $T = 6, C = 58, A = 70, G = 78$ | [58 78 70 6] |
| 5 | EIIP (electron-ion interaction potential) | $T = 0.1335, C = 0.1340, A = 0.1260, G = 0.0806$ | [0.1340 0.8060 0.1260 0.1335] |
| 6 | PP (purine/pyrimidine) | $T/C = 1, A/G = -1$ | [1 -1 -1 1] |
| 7 | Paired numeric | $T/A = 1, C/G = -1$ | [-1 -1 1 1] |
| 8 | Nearest-neighbor based doublet | 0 – 15 for all possible doublets | [14 8 1 7] |
| 9 | Codon | 0 – 63 for all possible 64 Codons | [2 35 22 44] |
| 10 | Just-A | $A = 1, \text{rest} = 0$ | [0 0 1 0] |
| 11 | Just-C | $C = 1, \text{rest} = 0$ | [1 0 0 0] |
| 12 | Just-G | $G = 1, \text{rest} = 0$ | [0 1 0 0] |
| 13 | Just-T | $T = 1, \text{rest} = 0$ | [0 0 0 1] |

Numerical representations of DNA sequences analyzed for usability in genomic classification with ML-DSP. The second column lists the numerical representation name, the third column describes the rule it uses, and the fourth is the output of this rule for the input DNA sequence $S_1 = CGAT$ . For the nearest-neighbor based doublet representation and codon representation, the DNA sequence is considered to be wrapped (the last position is followed by the first).

The datasets used for this analysis were the same as those in Table 1. The supervised machine learning classifiers used for this analysis were the six classifiers listed in the Methods section, with the exception of the datasets with more than 2,000 sequences where two of the classifiers (Subspace Discriminant and Subspace KNN) were omitted as being too slow. The results and the average accuracy scores for all these numerical representations, classifiers and datasets are summarized in Table 3.

**Table 3.**
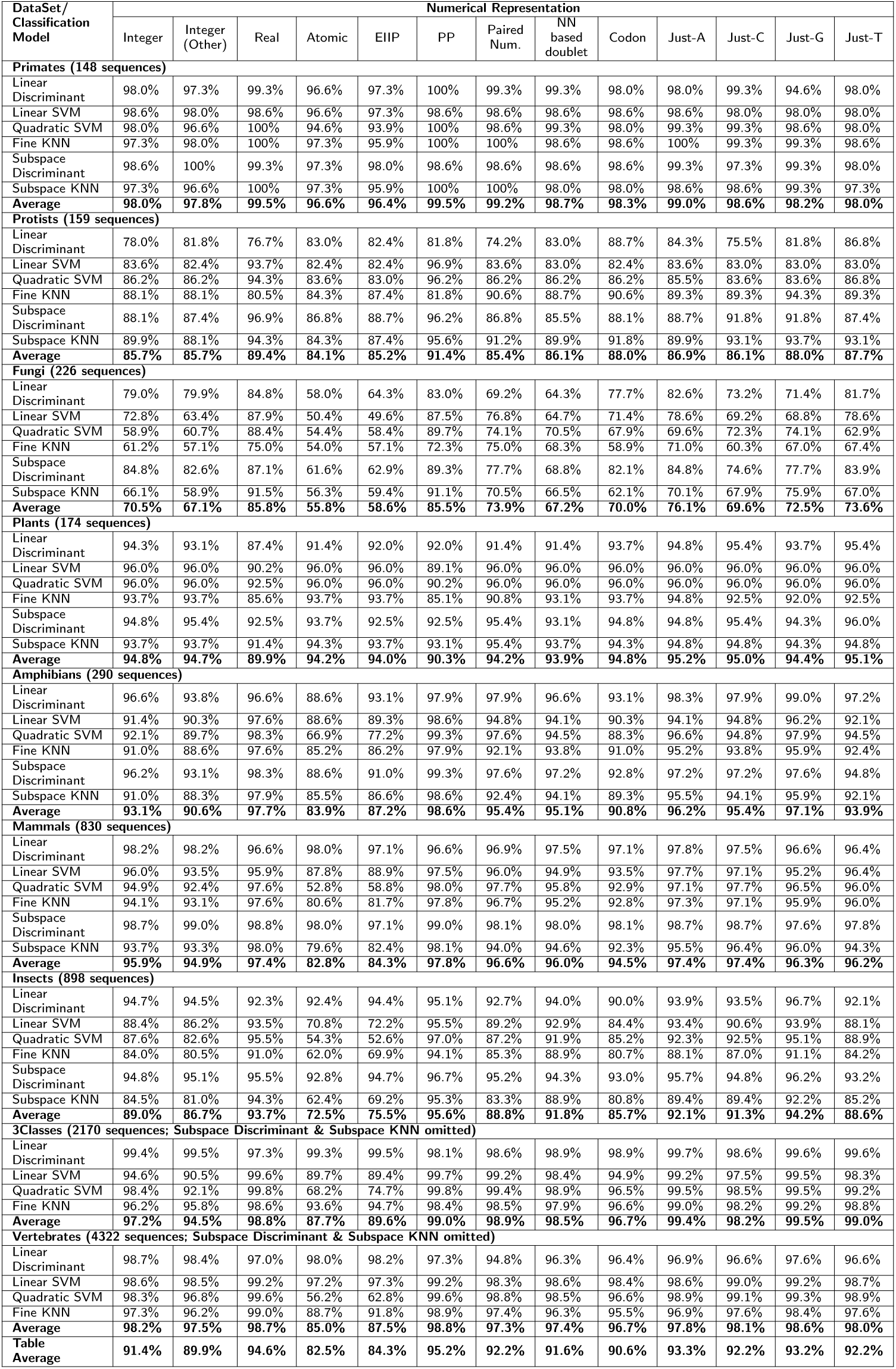
Average classification accuracies for 13 numerical representations.

As can be observed from Table 3, for all numerical representations, the table average accuracy scores (last row: average of averages, first over the six classifiers for each dataset, and then over all datasets), are high. Surprisingly, even using a single nucleotide numerical representation, which treats three of the nucleotides as being the same, and singles out only one of them (Just-A), results in an average accuracy of 93.3%. The best accuracy, for these datasets, is achieved when using the Purine/Pyrimidine (PP) representation, which yields an average accuracy of 95.2%.

For several of the numerical representations the average classification accuracies are so close that one could not choose a clear winner. For subsequent experiments we selected the top three representations in terms of accuracy scores: PP, Real and Just-A numerical representations.

### ML-DSP for three classes of vertebrates

As an application of ML-DSP using the Purine/Pyrimidine numerical representation for DNA sequences, we analyzed the set of vertebrate mtDNA genomes (median length 16,606 bp). The MoDMap, i.e., the multi-dimensional scaling 3D visualization of the genome interrelationships as described by the distances in the distance matrix, is illustrated in Fig 1. The dataset contains 3,740 complete mtDNA genomes: 553 bird genomes, 2,313 fish genomes, and 874 mammalian genomes. Quantitatively, the classification accuracy score obtained by the Quadratic SVM classifier was 100%.

**Figure 1.**
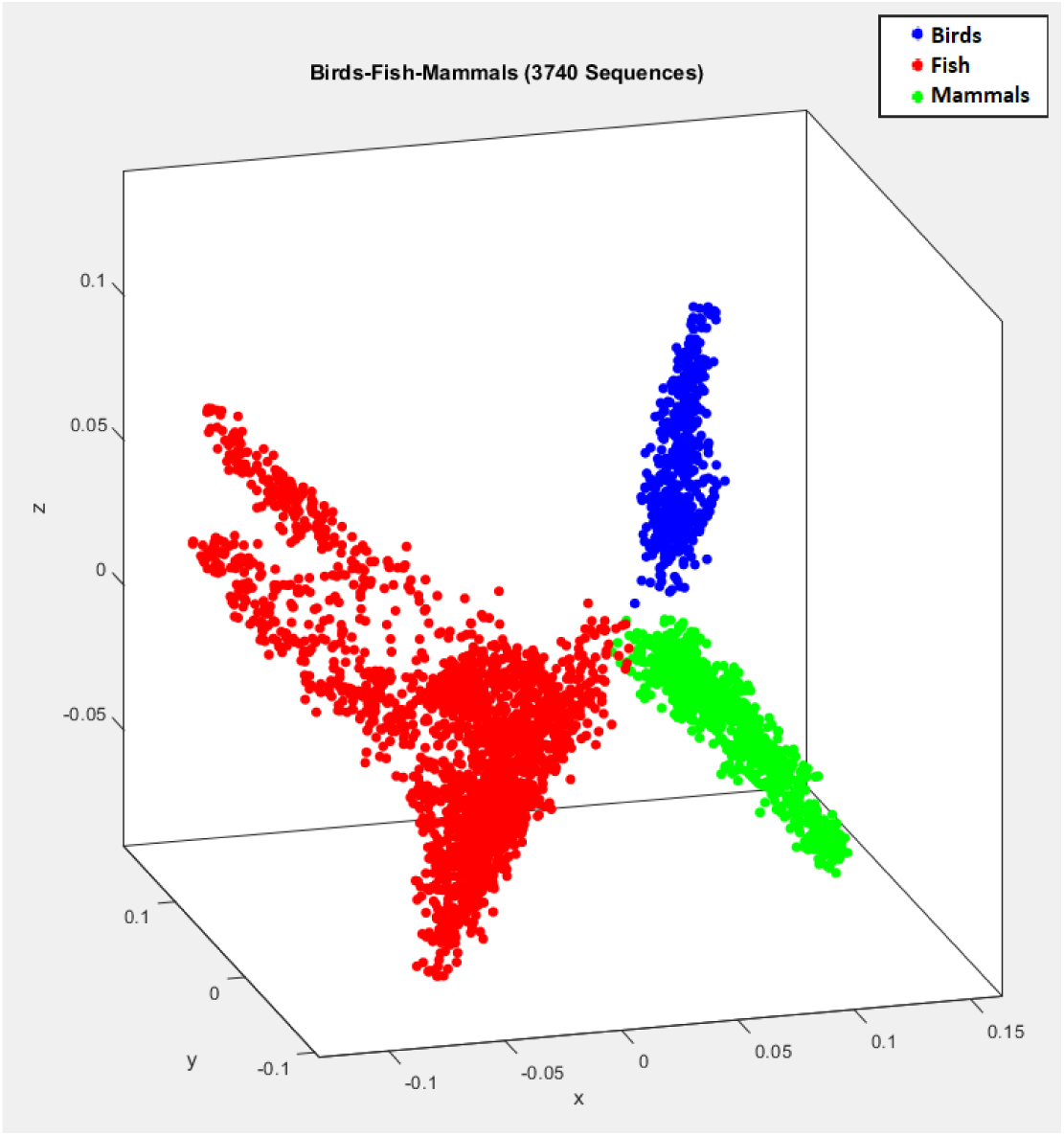
MoDMap of 3,740 full mtDNA genomes in subphylum Vertebrata, into three classes: Birds (blue, Aves: 553 genomes), fish (red, Actinopterygii 2,176 genomes, Chondrichthyes 130 genomes, Coelacanthiformes 2 genomes, Dipnoi 5 genomes), and mammals (green, Mammalia: 874 genomes). The accuracy of the ML-DSP classification into three classes, using the Quadratic SVM classifier, with the PP numerical representation, and PCC between magnitude spectra of DFT, was 100%.

### Classifying genomes with ML-DSP, at all taxonomic levels

We tested the ability of ML-DSP to classify complete mtDNA sequences at various taxonomic levels. For every dataset, we tested using the PP, Real, and Just-A numerical representations.

The starting point was domain Eukaryota (7, 396 sequences), which was classified into kingdoms, then kingdom Animalia was classified into phyla, etc. At each level, we picked the cluster with the highest number of sequences and then classified it into the next taxonomic level sub-clusters. The lowest level classified was family Cyprinidae (81 sequences) into its six genera. The maximum classification accuracy scores among the six classifiers, for the different taxonomic levels are shown in the Table 4.

**Table 4.**
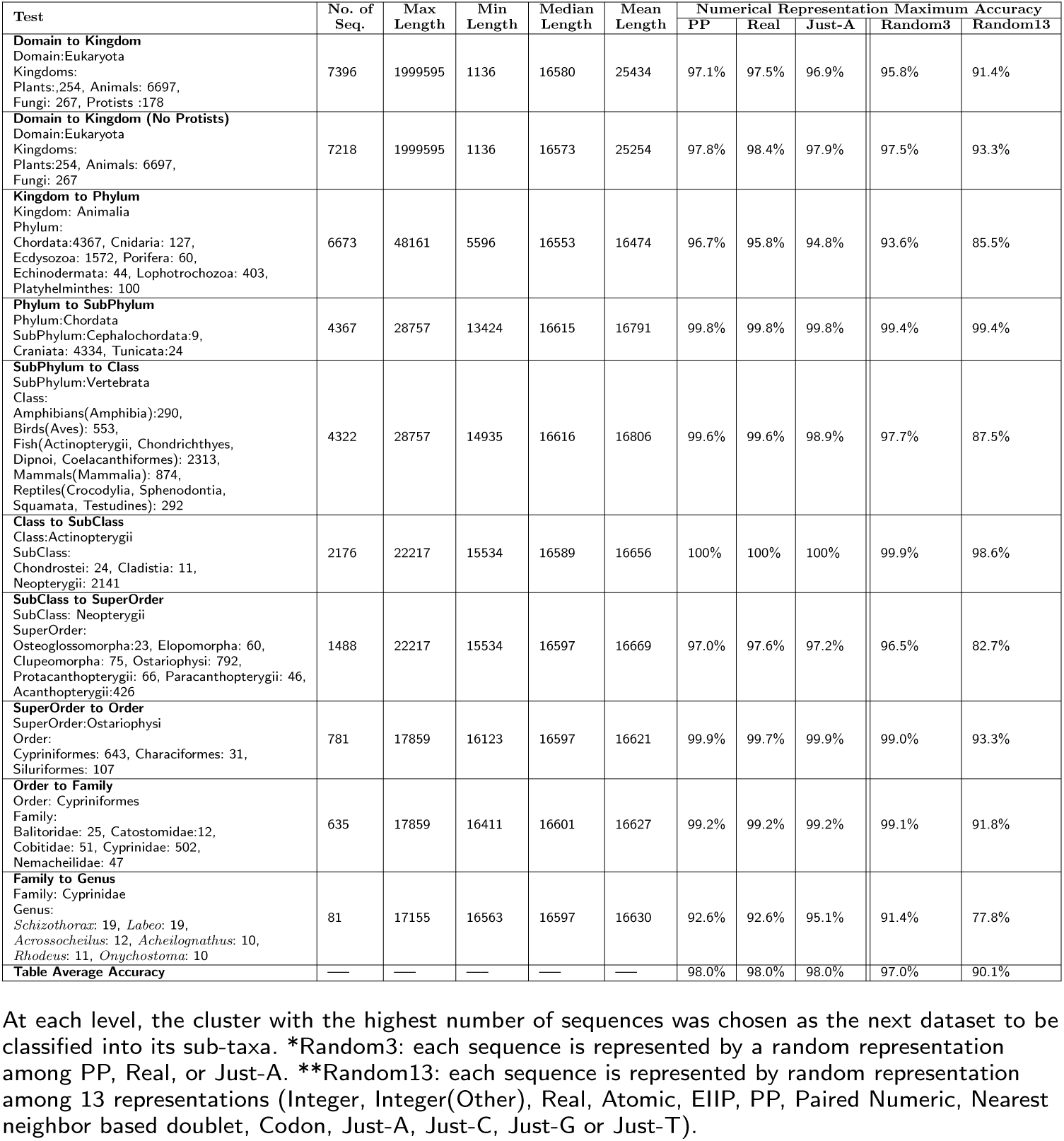
Maximum classification accuracy of ML-DSP (among the six classifiers), at different taxonomic levels.

Note that, at each taxonomic level, the maximum classification accuracy scores (among the six classifiers) for each of the three numerical representations considered are high, ranging from 92.6% to 100%, with only three scores under 95%. As this analysis also did not reveal a clear winner among the top three numerical representations, the question then arose whether the numerical representation we use mattered at all. To answer this question, we performed two additional experiments, that exploit the fact that the Pearson correlation coefficient is scale independent, and only looks for a pattern while comparing signals. For the first experiment we selected the top three numerical representations (PP, Real, Just-A) and, for each sequence in a given dataset, a numerical representation among these three was randomly chosen, with equal probability, to be the digital signal that represents it. The results are shown under the column “Random3” in Table 4: The maximum accuracy score over all the datasets is 97%. This is almost the same as the accuracy obtained when one particular numerical representation was used (1% lower, which is well within experimental error). We then repeated this experiment, this time picking randomly from any of the thirteen numerical representations considered. The results are shown under the column “Random13” in Table 4, with the table average accuracy score being 90.1%.

Overall, our results suggest that all three numerical representations PP, Real and Just-A have very high classifications accuracy scores (average 98%), and even a random choice of one of these representations for each sequence in the dataset does not significantly affect the classification accuracy score of ML-DSP (average 97%).

We also note that, in addition to being highly accurate in its classifications, MLDSP is ultrafast. Indeed, even for the largest dataset in Table 1, subphylum Vertebrata (4,322 complete mtDNA genomes, average length 16,806 bp), the distance matrix computation (which is the bulk of the classification computation) lasted under 5 seconds. Classifying a new primate mtDNA genome took 0.06 seconds when trained on 148 primate mtDNA genomes, and classifying a new vertebrate mtDNA genome took 7 seconds when trained on the 4,322 vertebrate mtDNA genomes.

### MoDMap visualization vs. ML-DSP quantitative classification results

The hypothesis tested by the next experiments was that the quantitative accuracy of the classification of DNA sequences by ML-DSP would be significantly higher than suggested by the visual clustering of taxa in the MoDMap produced with the same pairwise distance matrix.

As an example, the MoDMap in Fig 2(A), visualizes the distance matrix of mtDNA genomes from family Cyprinidae (81 genomes) with its genera *Acheilognathus* (10 genomes), *Rhodeus* (11 genomes), *Schizothorax* (19 genomes), *Labeo* (19 genomes), *Acrossocheilus* (12 genomes), *Onychostoma* (10 genomes); only the genera with at least 10 genomes are considered. The MoDMap seems to indicate an overlap between the clusters *Acheilognathus* and *Rhodeus*, which is biologically plausible as these genera belong to the same sub-family Acheilognathinae. However, when zooming in by plotting a MoDMap of only these two genera, as shown in Fig 2(B), one can see that the clusters are clearly separated visually. This separation is confirmed by the fact that the accuracy score of the Quadratic SVM classifier for the dataset in Fig 2(B) is 100%. The same quantitative accuracy score for the classification of the dataset in Fig 2(A) with Quadratic SVM is 92.6%, which intuitively is much better than the corresponding MoDMap would suggest. This is likely due to the fact that the MoDMap is a three-dimensional approximation of the positions of the genome-representing points in a multi-dimensional space (the number of dimensions is (*n −* 1), where *n* is the number of sequences).

**Figure 2.**
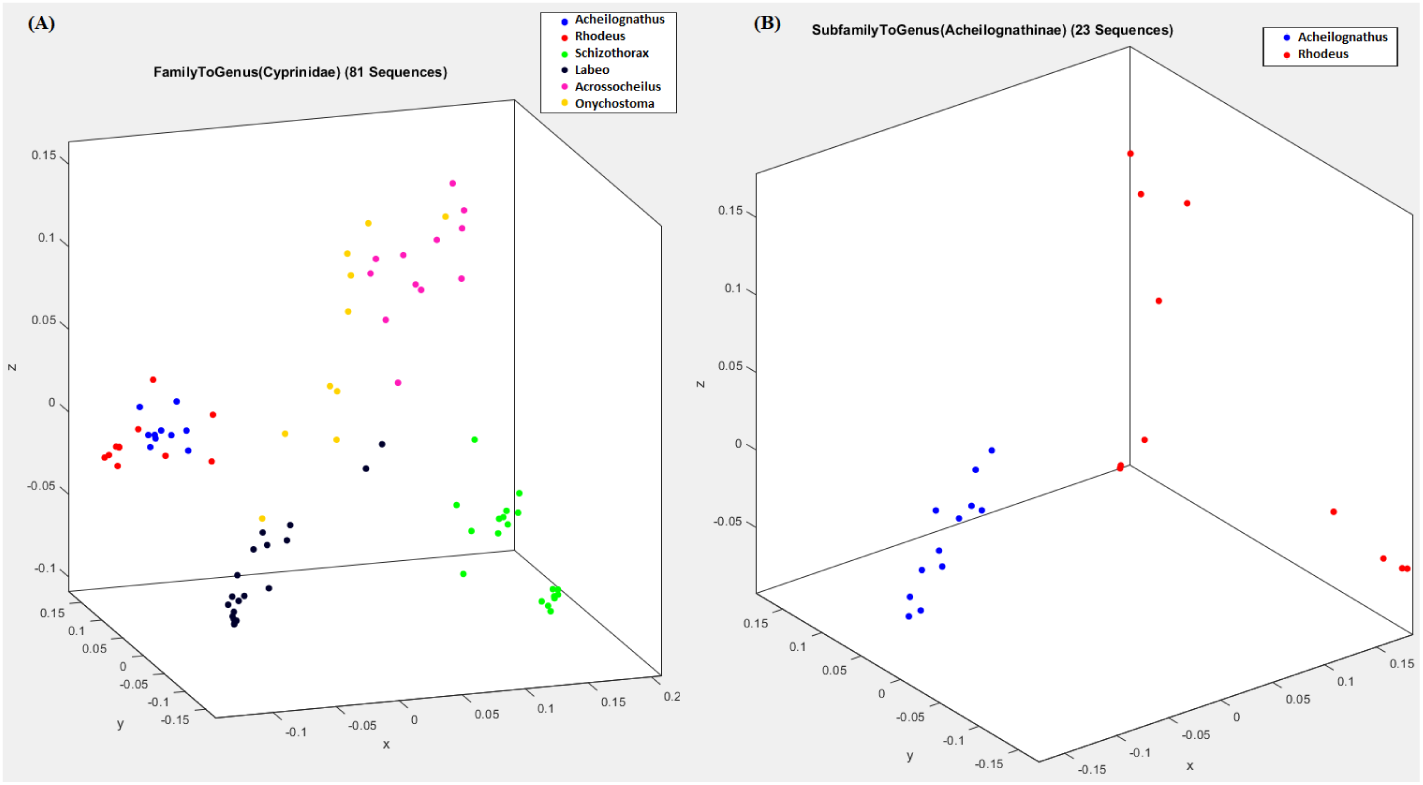
MoDMap of family Cyprinidae and its genera. (A): Genera *Acheilognathus* (blue, 10 genomes), *Rhodeus* (red, 11 genomes), *Schizothorax* (green, 19 genomes), *Labeo* (black, 19 genomes), *Acrossocheilus* (magenta, 12 genomes), *Onychostoma* (yellow, 10 genomes); (B): Genera *Acheilognathus* and *Rhodeus*, which overlapped in (A), are visually separated when plotted separately in (B). The classification accuracy with Quadratic SVM of the dataset in (A) was 92.6%, and of the dataset in (B) was 100%.

This being said, MoDMaps can still serve for exploratory purposes. For example, the MoDMap in Fig 2(A) suggests that species of the genus *Onychostoma* (subfamily listed “unknown” in NCBI) (yellow), may be genetically related to species of the genus *Acrossocheilus* (subfamily Barbinae) (magenta). Upon further exploration of the distance matrix, one finds that indeed the distance between the centroids of these two clusters is lower than the distance between each of these two cluster-centroids to the other cluster-centroids. This supports the hypotheses, based on morphological evidence [46], that genus *Onychostoma* belongs to the subfamily Barbinae, respectively that genus *Onychostoma* and genus *Acrossocheilus* are closely related [47]. Note that this exploration, suggested by MoDMap and confirmed by calculations based on the distance matrix, could not have been initiated based on ML-DSP alone (or other supervised machine learning algorithms), as ML-DSP only predicts the classification of new genomes into one of the taxa that it was trained on, and does not provide any other additional information.

As another comparison point between MoDMaps and supervised machine learning outputs, Fig 3(A) shows the MoDMap of the superorder Ostariophysi with its orders Cypriniformes (643 genomes), Characiformes (31 genomes) and Siluriformes (107 genomes). The MoDMap shows the clusters as overlapping, but the Quadratic SVM classifier that quantitatively classifies these genomes has an accuracy of 99.9%. Indeed, the confusion matrix^[1]^ in Fig 3(B) shows that Quadratic SVM mis-classifies only 1 sequence out of 781. This indicates that when the visual representation in a MoDMap shows cluster overlaps, this may only be due to the dimensionality reduction to three dimensions, while ML-DSP actually provides a much better quantitative classification based on the same distance matrix.

**Figure 3.**
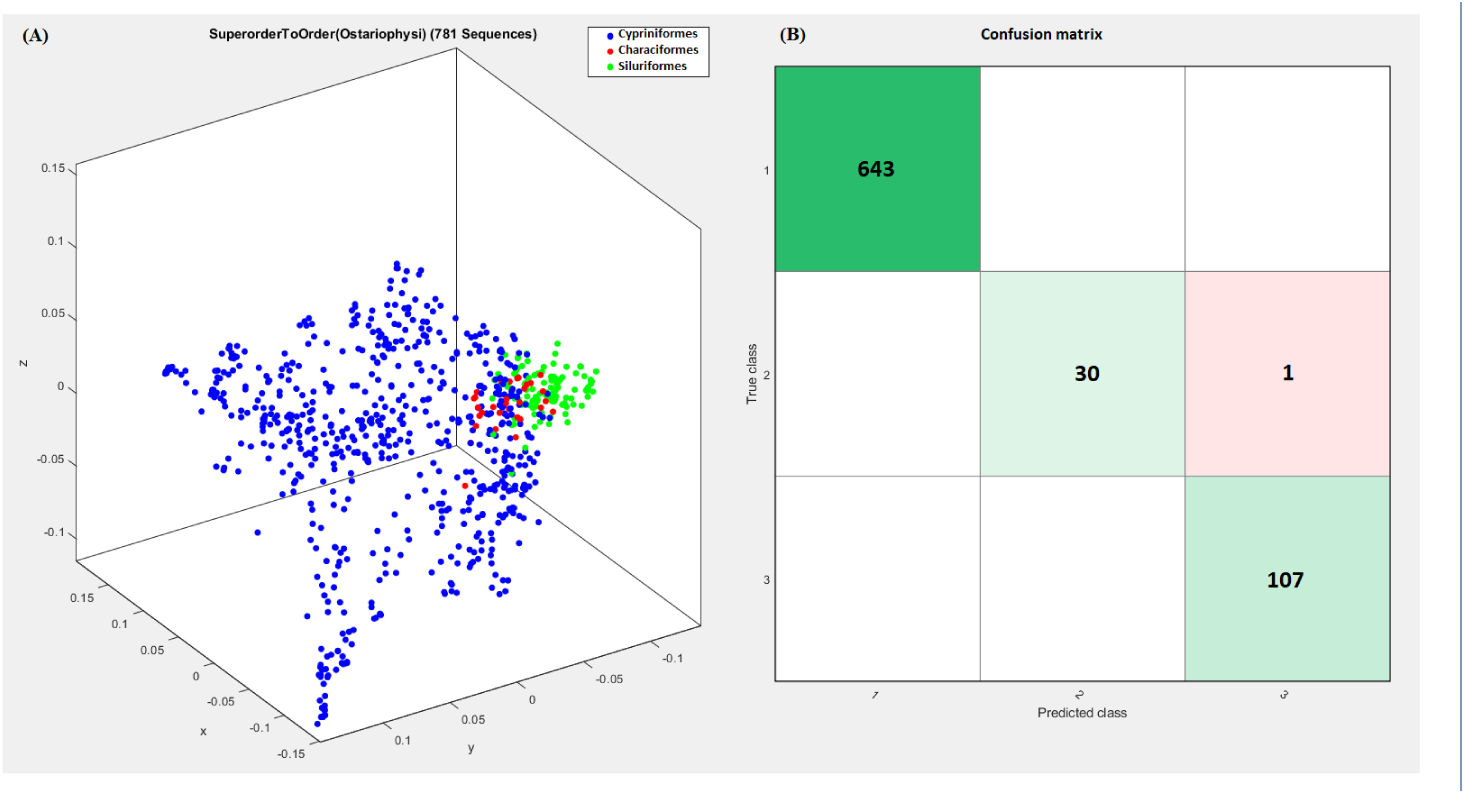
MoDMap of the superorder Ostariophysi, and the confusion matrix for the Quadratic SVM classification of this superorder into orders. (A): MoDMap of orders Cypriniformes (blue, 643 genomes), Characiformes (red, 31 genomes), Siluriformes (green, 107 genomes). (B): The confusion matrix generated by Quadratic SVM, illustrating its true class vs. predicted class performance (top-to-bottom and left-to-right: Cypriniformes, Characiformes, Siluriformes). The numbers in the squares on the top-left to bottom-right diagonal (green) indicate the numbers of correctly classified DNA sequences, by order. The off-diagonal red square indicates that 1 mtDNA genome of the order Characiformes has been erroneously predicted to belong to the order Siluriformes. The Quadratic SVM that generated this confusion matrix had a 99.9% classification accuracy.

### Applications to other genomic datasets: dengue virus subtyping

The experiments in this section indicate that the applicability of our method is not limited to mitochondrial DNA sequences. Fig 4 shows the MoDMap of all 4,721 complete dengue virus sequences available in NCBI on August 10, 2017, into the subtypes DENV-1 (2,008 genomes), DENV-2 (1,349 genomes), DENV-3 (1,010 genomes), DENV-4 (354 genomes). The average length of these complete viral genomes is 10,595 bp. Despite the dengue viral genomes being very similar, the classification accuracy of this dataset into subtypes, using the Quadratic SVM classifier, was 100%.

**Figure 4.**
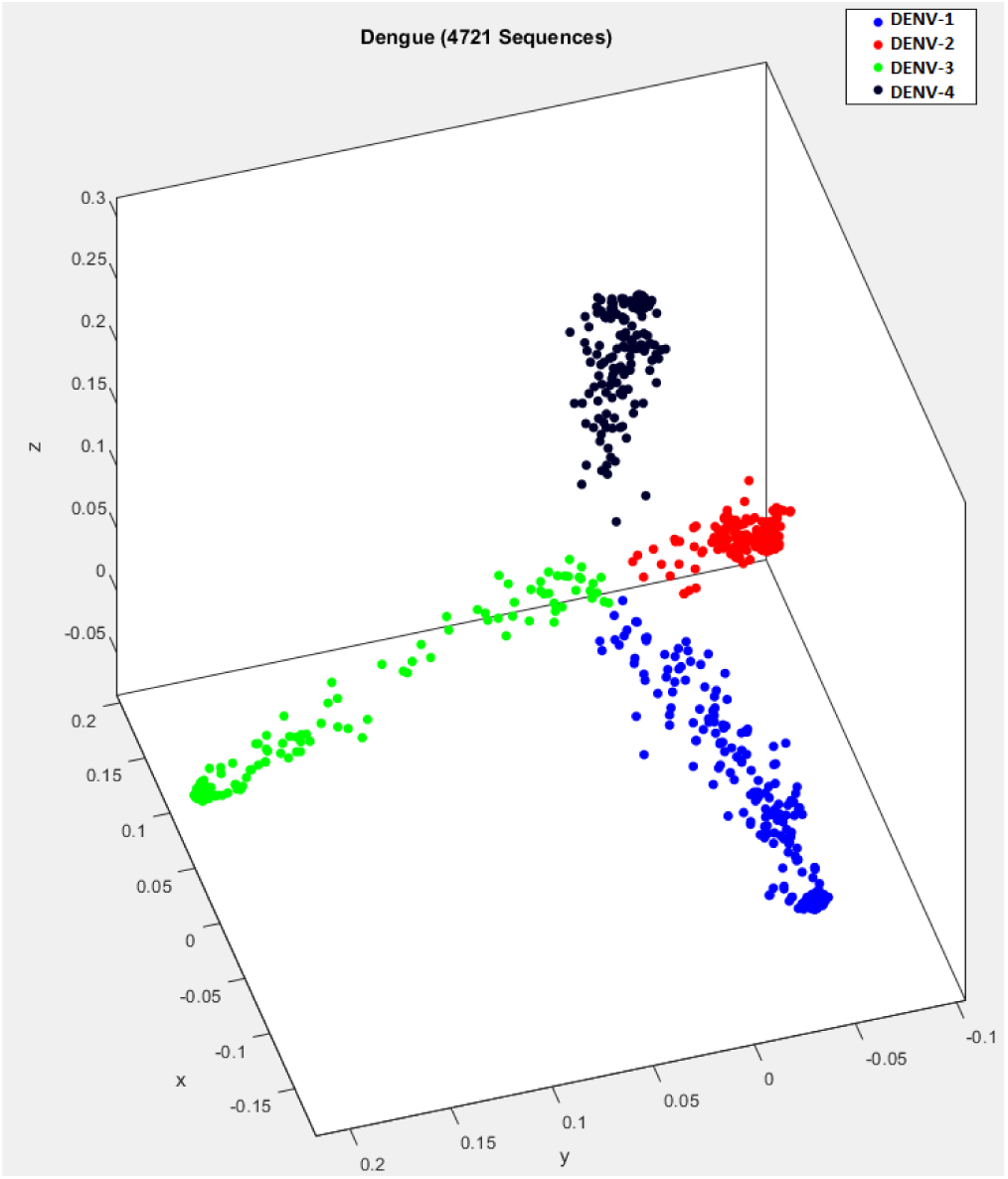
MoDMap of 4,271 dengue virus genomes and their subtypes. The colours represent virus subtypes DENV-1 (blue, 2008 genomes), DENV-2 (red, 1349 genomes), DENV-3 (green, 1010 genomes), DENV-4 (black, 354 genomes); The classification accuracy of the Quadratic SVM classifier for this dataset was 100%.

### Comparison of ML-DSP with state-of-the-art alignment-based and alignment-free tools

The computational experiments in this section compare ML-DSP with three state-of-the-art alignment-based and alignment-free methods: the alignment-based tool MEGA7 [3] with alignment using MUSCLE [4] and CLUSTALW [5, 6], and the alignment-free method FFP (Feature Frequency Profiles) [28].

For this performance analysis we selected three datasets. The first two datasets are benchmark datasets used in other genetic sequence comparison studies [44]: The first dataset comprises 38 influenza viral genomes, and the second dataset comprises 41 mammalian complete mtDNA sequences. The third datase, of our choice, is much larger, consisting of 4, 322 vertebrate complete mtDNA sequences, and was selected to compare scalability.

For the alignment-based methods, we used the distance matrix calculated in MEGA7 from sequences aligned with either MUSCLE or CLUSTALW. For the alignment-free FFP, we used the default value of *k* = 5 for *k*-mers (a *k*-mer is any DNA sequence of length *k*; any increase in the value of the parameter *k*, for the first dataset, resulted in a lower classification accuracy score for FFP). For ML-DSP we chose the Integer numerical representation and computed the average classification accuracy over all six classifiers for the first two datasets, and over all classifiers except Subspace Discriminant and Subspace KNN for the third dataset.

Table 5 shows the performance comparison (classification accuracy and processing time) of these four methods. The processing time included all computations, starting from reading the datasets to the completion of the distance matrix - the common element of all four methods. The listed processing times do not include the time needed for the computation of phylogenetic trees, MoDMap visualizations, or classification.

**Table 5.** Comparison of classification accuracy and processing time for the distance matrix computation with MEGA7(MUSCLE), MEGA7(CLUSTALW), FPP, and ML-DSP.

| DataSet | Parameter | MEGA7<br>(MUSCLE) | MEGA7<br>(CLUSTALW) | FFP | ML-DSP |
| --- | --- | --- | --- | --- | --- |
| **Influenza Virus**<br>(38 sequences)<br>Average Length: 1407bp | Maximum Classification Accuracy | 97.4% | 97.4% | 76.3% | 100% |
|  | Average Classification Accuracy | 95.2% | 95.6% | 64.0% | 97.4% |
|  | Processing Time | 7.44 sec | 2 min 14 sec | 0.2 sec | 0.2 sec |
| **Mammalia**<br>(41 sequences)<br>Average Length: 16647bp | Maximum Classification Accuracy | 95.1% | 92.7% | 61.0% | 92.7% |
|  | Average Classification Accuracy | 93.2% | 92.7% | 52.2% | 89.2% |
|  | Processing Time | 11 min 15sec | 5 hr 38 min | 0.3 sec | 0.3 sec |
| **Vertebrates**<br>(4322 sequences)<br>Average Length: 16806bp | Maximum Classification Accuracy | — | — | 99.1% | 99.5% |
|  | Average Classification Accuracy | — | — | 97.0% | 98.7% |
|  | Processing Time | >2 hours | >6 hours | 94 sec | 28 sec |

As seen in Table 5, ML-DSP outperforms FFP in accuracy for all three testing datasets. In terms of processing time, ML-DSP and FFP perform similarly for the two small datasets, but for the largest dataset ML-DSP is three times faster, which indicates that ML-DSP is significantly more scalable to larger datasets.

The alignment-based tool MEGA7(MUSCLE) had similar classification accuracy scores as ML-DSP for the first dataset, but was much slower. For the second dataset, MEGA7(MUSCLE) obtained average accuracy scores 4% higher than ML-DSP^[2]^, at the cost of being *≈* 2, 250 times slower. MEGA7(CLUSTALW) had similar classification accuracy as MEGA7(MUSCLE) while being much slower, and 67,600 times slower than ML-DSP. For the third and largest dataset, neither MEGA7(MUSCLE) nor MEGA7(CLUSTALW) could complete the alignment after running for 2 hours and 6 hours respectively, and had to be terminated, while ML-DSP only took 28 seconds.

Overall, this comparison indicates that, for these datasets, the alignment-free methods (ML-DSP and FFP) show a clear advantage over the alignment-based methods MEGA7(MUSCLE) and MEGA7(CLUSTALW)) in terms of processing time. Among the two alignment-free methods compared, ML-DSP outperforms FFP in classification accuracy, for both the benchmark datasets as well as the larger vertebrates dataset.

As another angle of comparison, Fig 5 displays the MoDMaps of the first benchmark dataset (38 influenza virus genomes) produced from the distance matrices generated by FFP, MEGA7(MUSCLE), MEGA7(CLUSTALW), and ML-DSP respectively. Fig 5(A) shows that with FFP it is difficult to observe any visual separation of the dataset into subtype clusters. Fig 5(B), MEGA7(MUSCLE), and Fig 5(C) MEGA7(CLUSTALW) show overlaps of the clusters of points represent-ing subtypes H1N1 and H2N2. In contrast, Fig 5(D), which visualizes the distance matrix produced by ML-DSP, shows a clear separation among all subtypes.

**Figure 5.**
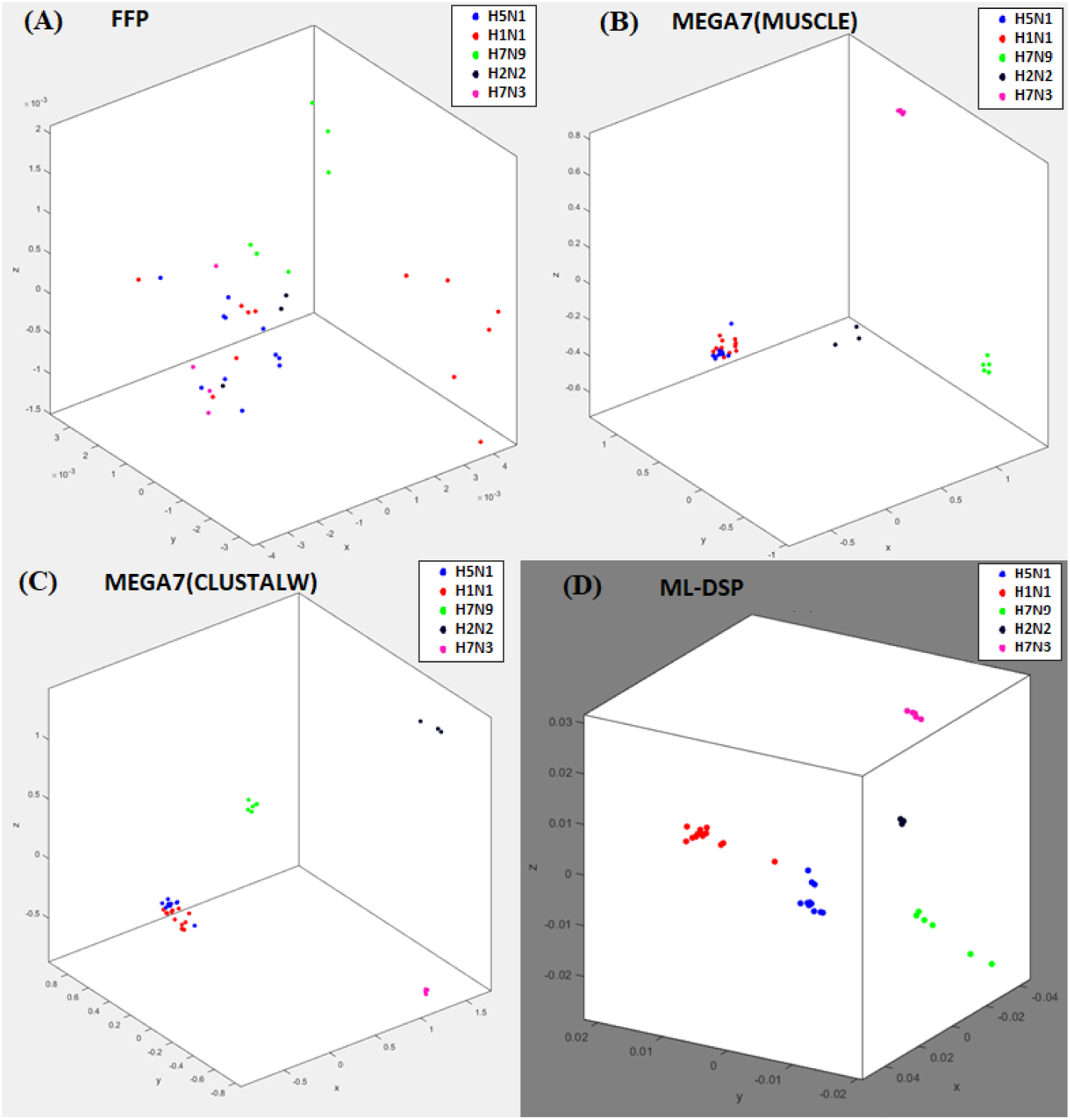
MoDMaps of the influenza virus dataset from Table 5, based on the four methods. The points represent viral genomes of subtypes H1N1 (red, 13 genomes), H2N2 (black, 3 genomes), H5N1 (blue, 11 genomes), H7N3 (magenta, 5 genomes), H7N9 (green, 6 genomes); ModMaps are generated using distance matrices computed with (A) FFP; (B) MEGA7(MUSCLE); (C) MEGA7(CLUSTALW); (D) ML-DSP.

Finally Figures 6 and 7 display the phylogenetic trees generated by each of the four methods considered. Fig 6(A), the tree generated by FFP, has many misclassified genomes, which was expected given the MoDMap visualization of its distance matrix in Fig 5(A). Fig 7(A) displays the phylogenetic tree generated by MEGA7, which was the same for both MUSCLE and CLUSTALW: It has only one incorrectly classified H5N1 genome, placed in middle of H1N1 genomes. Fig 6(B) and Fig 7(B) display the phylogenetic tree generated using the distance produced by ML-DSP (shown twice, in parallel with the other trees, for ease of comparison). ML-DSP classified all genomes correctly.

**Figure 6.**
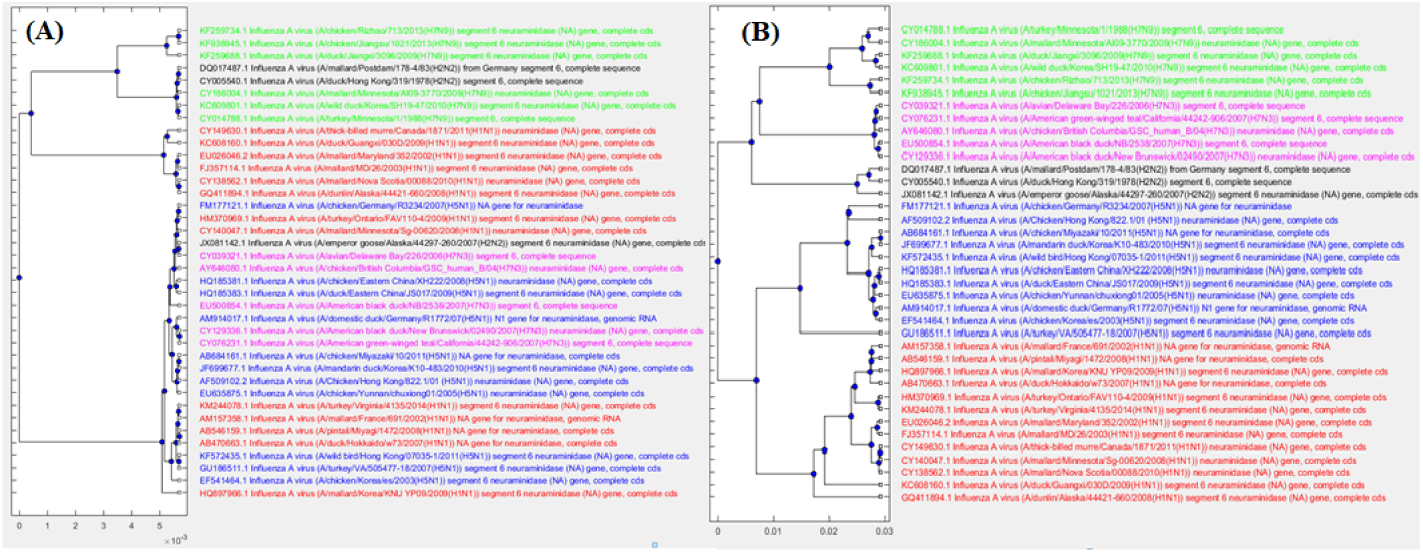
Phylogenetic tree comparison: FFP with ML-DSP. The phylogenetic tree generated for 38 influenza virus genomes using (A): FFP (B): ML-DSP.

**Figure 7.**
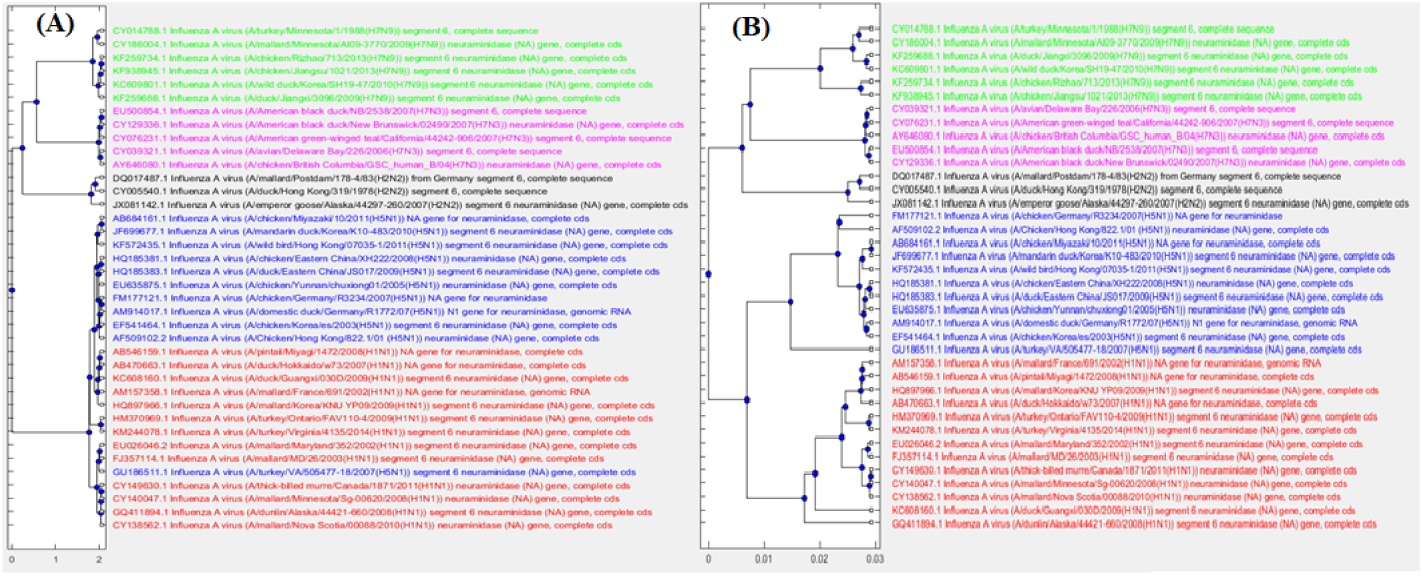
Phylogenetic tree comparison: MEGA7(MUSCLE/CLUSTALW) with ML-DSP. The phylogenetic tree generated for 38 influenza virus genomes using (A): MEGA7(MUSCLE/CLUSTALW) (B): ML-DSP.

## Discussion

The computational efficiency of ML-DSP is due to the fact that it is alignment-free (hence it does not need multiple sequence alignment), while the combination of 1D numerical representations, Discrete Fourier Transform and Pearson Correlation Coefficient makes it extremely computationally time efficient, and thus scalable.

ML-DSP is not without limitations. We anticipate that the need for equal length sequences and use of length normalization could introduce issues with examination of small fragments of larger genome sequences. Usually genomes vary in length and length normalization always results in adding (up-sampling) or losing (down-sampling) some information. Although the Pearson Correlation Coefficient can distinguish the signal patterns even in small sequence fragments, and we did not find any considerable disadvantage while considering complete mitochondrial DNA genomes with their inevitable length variations, length normalization may cause issues when we deal with the fragments of genomes, and the much larger nuclear genome sequences.

Lastly, ML-DSP has two drawbacks, inherent in any supervised machine learning algorithm. The first is that ML-DSP is a black-box method which, while producing a highly accurate classification prediction, does not offer a (biological) explanation for its output. The second is that it relies on the existence of a training set from which it draws its “knowledge”, that is, a set consisting of known genomic sequences and their taxonomic labels. ML-DSP uses such a training set to “learn” how to classify new sequences into one of the taxonomic classes that it was trained on, but it is not able to assign it to a taxon that it has not been exposed to.

## Conclusions

We proposed ML-DSP, an ultrafast and accurate alignment-free supervised machine learning classification method based on digital signal processing of DNA sequences (and its software implementation). ML-DSP successfully addresses the limitations of alignment-free methods identified in [7], as follows:

i. `Lack of software implementation:` ML-DSP is supplemented with freely available source-code. The ML-DSP software can be used with the provided datasets or any other custom dataset and provides the user with any or all of: pairwise distances, 3D sequence interrelationship visualization, phylogenetic trees, or classification accuracy scores.
ii. `Use of simulated sequences or very small real-world datasets:` ML-DSP was successfully tested on a variety of real-world datasets. We considered all complete mitochondrial DNA sequences available on NCBI at the time of this study. Also, we tested ML-DSP in different evolutionary scenarios such as different levels of taxonomy (domain to genus), small dataset (38 sequences), large dataset (4,322 sequences), short sequences (1,136 bp), long sequences (1,999,595 bp), benchmark datasets of influenza virus and mammalian mtDNA genomes etc.
iii. `Memory overhead:` ML-DSP uses neither *k*-mers nor any compression algorithms. Thus, scalability does not cause an exponential memory overhead, and high accuracy is preserved on large datasets.

## Methods

The main idea behind ML-DSP is to combine supervised machine learning techniques with digital signal processing, for the purpose of DNA sequence classification. More precisely, for a given set *S* = {*S*_1_, *S*_2_,…, *S*_*n*_} of *n* DNA sequences, ML-DSP uses:

- `DNA numerical representations` to obtain a set *N* = {*N*_1_*, N*_2_*,…, N_n_*} where *N*_*i*_ is a discrete numerical representation of the sequence *S*_*i*_, 1 *≤ i ≤ n*.
- `Discrete Fourier Transform (DFT)` applied to the length-normalized digital signals *N*_*i*_, to obtain the frequency distribution; the magnitude spectrum *M*_*i*_ of this frequency distribution is then obtained.
- `Pearson Correlation Coefficient (PCC)` to compute the distance matrix of all pairwise distances for each pair of magnitude spectra (*M*_*i*_, *M*_*j*_), where 1 *≤ i, j ≤ n*.
- `Supervised Machine Learning` classifiers which take the pairwise distance matrix for a set of sequences, together with their respective taxonomic labels, in a training set, and output the taxonomic classification of a new DNA sequence. To measure the performance of such a classifier, we use the 10-fold crossvalidation technique.
- Independently, `Classical MultiDimensional Scaling (MDS)` takes the distance matrix as input and returns an (*n × q*) coordinate matrix, where *n* is the number of points (each point represents a unique sequence from set *S*) and *q* is the number of dimensions. The first three dimensions are used to display a MoDMap, which is the simultaneous visualization of all points in 3*D*-space.

### DNA numerical representations

To apply digital signal processing techniques to genomic data, genomic sequences are first mapped into discrete numerical representations of genomic sequences, called *genomic signals* [48].

In our analysis of various numerical representations for DNA sequences, we considered only 1*D* numerical representations, that is, those which produce a single output numerical sequence, called also *indicator sequence*, for a given input DNA sequence. We did not consider other numerical representations, such as binary [29], or nearest dissimilar nucleotide [49], because those generate four numerical sequences for each genomic sequence, and would thus not be scalable to classifications of thousands of complete genomes.

### Discrete Fourier Transform (DFT)

Our alignment-free classification method of DNA sequences makes use of the Discrete Fourier Transform (DFT) magnitude spectrum of the discrete numerical sequences (discrete digital signals) that represent DNA sequences. In some sense, these DFT magnitude spectra reflect the nucleotide distribution of the originating DNA sequences.

To start with, assuming that all input DNA sequences have the same length *p*, for each DNA sequence *S*_*i*_ = (*S*_*i*_(0)*, S_i_*(1),…, *S*_*i*_(*p −* 1)), in the input dataset, where 1 *≤ i ≤ n*, *S*_*i*_(*k*) *∈* {*A, C, G, T*}, 0 *≤ k ≤ p −* 1, we calculate its corresponding discrete numerical representation (discrete digital signal) *N*_*i*_ defined as

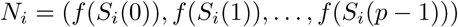

where, for each 0 *≤ k ≤ p−*1, the quantity *f* (*S*_*i*_(*k*)) is the value under the numerical representation *f* of the nucleotide in the position *k* of the DNA sequence *S*_*i*_.

Then, the DFT of the signal *N*_*i*_ is computed as the vector *F*_*i*_ where, for 0 *≤ k ≤ p −* 1 we have

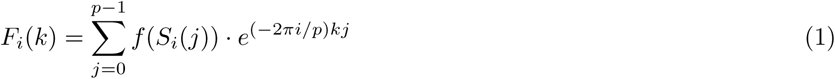

The magnitude vector corresponding to the signal *N*_*i*_ can now be defined as the vector *M*_*i*_ where, for each 0 *≤ k ≤ p −* 1, the value *M*_*i*_(*k*) is the absolute value of *F*_*i*_(*k*), that is, *M*_*i*_(*k*) = *| F_i_*(*k*)*|*. The magnitude vector *M*_*i*_ is also called the magnitude spectrum of the digital signal *N*_*i*_ and, by extension, of the DNA sequence *S*_*i*_. For example, if the numerical representation *f* is Integer (row 1 in Table 2), then for the sequence *S*_1_ = *CGAT*, the corresponding numerical representation is *N*_1_= (1, 3, 2, 0), the result of applying DFT is *F*_1_ = (6*, −*1 − 3*i,* 0*, −*1 + 3*i*) and its magnitude spectrum is *M*_1_ = (6, 3.1623, 0, 3.1623).

Fig 8A shows the discrete digital signal (using the PP numerical representation, row 6 of Table 2) of the DNA sequence consisting of the first 100 bp of the mtDNA genome of *Branta canadensis* (Canada goose, NCBI accession number *NC* 007011.1), and of the DNA sequence consisting of the first 100 bp of the mtDNA genome of *Castor fiber* (European beaver; NCBI accession number *NC* 028625.1). Fig 8B shows the DFT magnitude spectra of the same two signals/sequences. As can be seen in Fig 8B, these mtDNA sequences exhibit different DFT magnitude spectrum patterns, and this can be used to distinguish them computationally by using. e.g., the Pearson correlation coefficient, as described in the next subsection. Other techniques have also been used for genome similarity analysis, for example comparing the phase spectra of the DFT of digital signals of full mtDNA genomes, as seen in Figure 9 and [50, 51].

**Figure 9.**
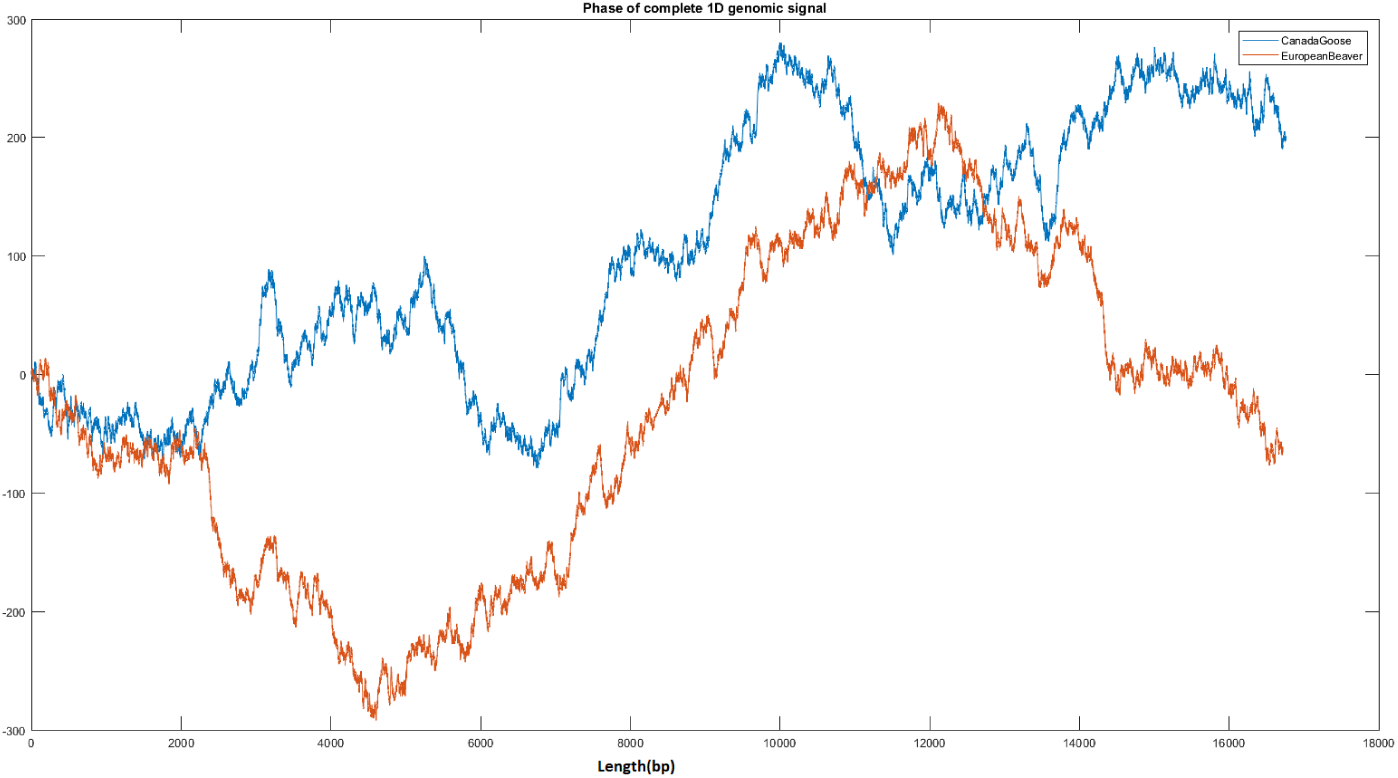
Canada goose (blue, 16,760 bp) vs. European beaver (red, 16,722 bp) - comparison between the DFT phase spectra of their full mtDNA genomes.

**Figure 8.**
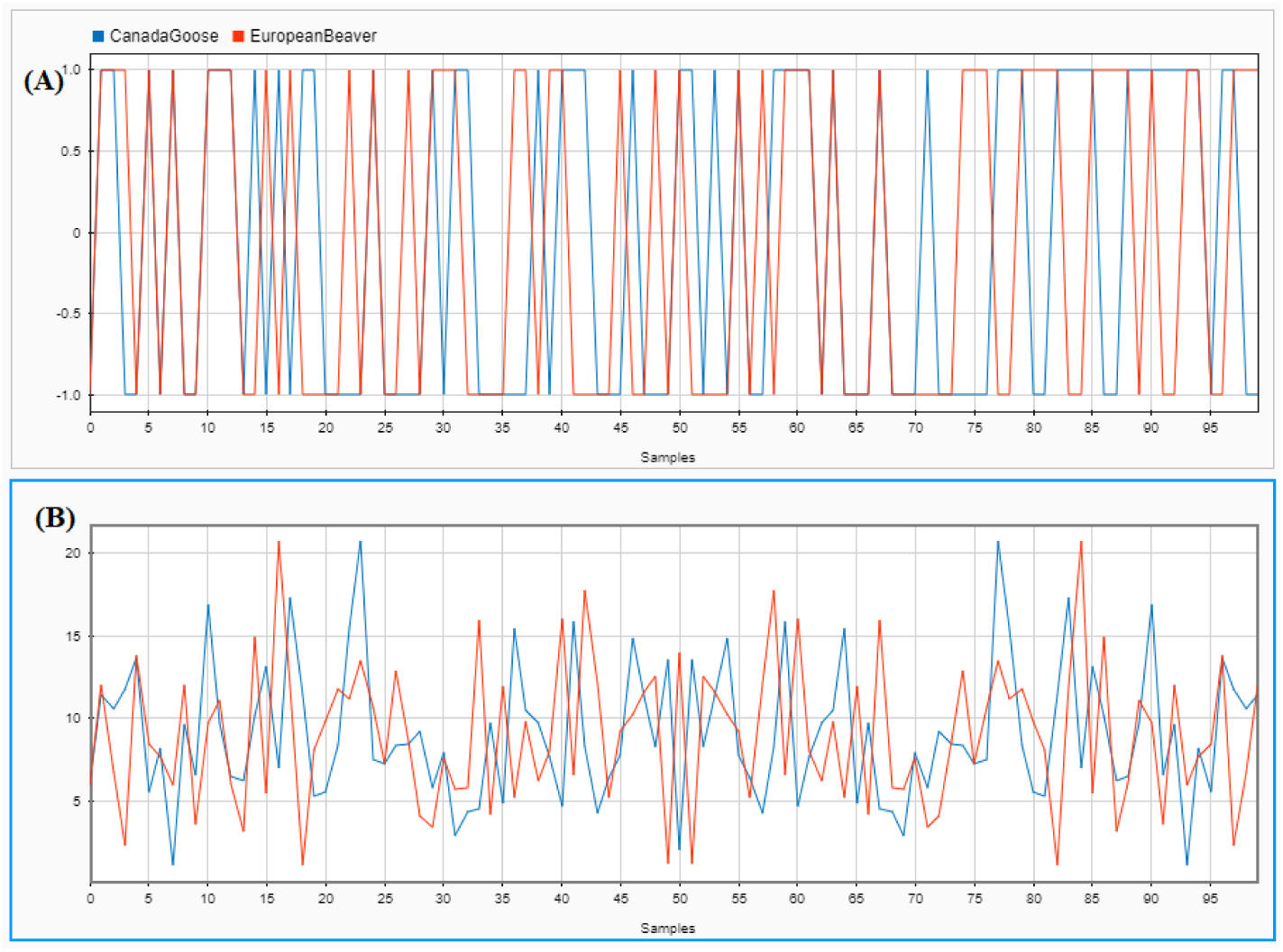
Canada goose (blue) vs European beaver (red): comparison of the DFT magnitude spectra of the first 100 bp of their mtDNA genomes. (A): Graphical illustration of the discrete digital signals of the respective DNA sequences, obtained using the PP representation. (B): DFT magnitude spectra of the signals in (A).

Note that, with the exception of the example in Fig 8, all of the computational experiments in this paper use full genomes.

### Pearson Correlation Coefficient (PCC)

Consider two variables *X* and *Y* (here *X* and *Y* are the magnitude spectra *M*_*i*_ and *M*_*j*_ of two signals), each of length *p*, that is, *X* = {*X*_0_*,…, X_p−1_*} and *Y* = {*Y*_0_*,…, Y_p-1_*}. The Pearson correlation coefficient *r*_*XY*_ between *X* and *Y* is the ratio of their covariance (measure of how much *X* and *Y* vary together) to the product of their standard deviations [52, 53], that is,

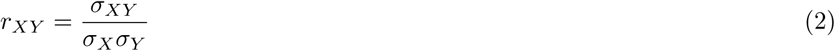

where the covariance of *X* and *Y* is 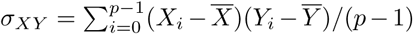, and the standard deviation is 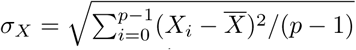, and similarly for *σ_Y_*, where the average is defined as 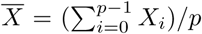 and similarly for *Y*. Now the formula for the Pearson Correlation Coefficient becomes

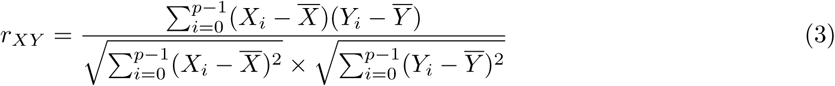

The Pearson correlation coefficient between *X* and *Y* is a measure of their linear correlation, and has a value between +1 (total positive linear correlation) and −1 (total negative linear correlation); 0 is no linear correlation. We normalized the results, by taking (1 − *r_XY_*)*/*2, to obtain distance values between 0 and 1 (value 0 for identical signals, and 1 for negatively correlated signals). For our data sets, the PCC values between any two digital signals of DNA sequences ranged between 0 and 0.6.

For each pairwise distance calculation, the Pearson correlation coefficient requires the input variables (that is, the magnitude spectra of the two sequences to be correlated) to have the same length. The length of a magnitude spectrum is equal to the length of corresponding numerical digital signal, which in turn is equal to the length of the originating DNA sequence. Given that genome sequences are typically of different lengths, it follows that their corresponding digital signals need to be length-normalized, if we are to be able to use the Pearson correlation coefficient. Hoang et al. avoided normalization and considered only the first few mathematical moments constructed from the power spectra for comparison, after applying DFT [54]. The limitation of this method is that one loses information that may be necessary for a meaningful comparison. This is specially important when the genomes compared are very similar to each other.

Different methods for length-normalizing digital signals were tested: down-sampling [55], up-sampling to the maximum length using zero padding [15], even scaling extension [56], periodic extension, symmetric padding, or anti-symmetric padding [57]. For example, zero-padding, which adds zeroes to all of the sequences shorter than the maximum length, was used in [30], e.g., for taxonomic classifications of ribosomal S18 subunit genes from twelve organisms. While this method may work for datasets of sequences of similar lengths, it is not suitable for datasets of sequences of very different lengths (our study: fungi mtDNA genomes dataset − 1,364 bp to 235,849 bp; plant mtDNA genomes dataset − 12,998 bp to 1,999,595 bp; protist mtDNA genomes dataset − 5,882 bp to 77,356 bp). In such cases, zero-padding acts as a tag and may lead to inadvertent classification of sequences based on their length rather than based on their sequence composition. Thus, we employed instead anti-symmetric padding, whereby, starting from the last position of the signal, boundary values are replicated in an anti-symmetric manner. We also considered two possible ways of employing anti-symmetric padding: normalization to the maximum length (where shorter sequences are extended to the maximum sequence length by anti-symmetric padding) vs. normalization to the median length (where shorter sequences are extended by anti-symmetric padding to the median length, while longer sequences are truncated after the median length).

### Supervised Machine Learning

In this paper we used the Linear discriminant, Linear SVM, Quadratic SVM, Fine KNN, Subspace discriminant and Subspace KNN classifiers from the Classification Learner application of MATLAB (Statistics and Machine Learning Toolbox).

To assess the performance of the classifiers, we used 10-fold cross validation. In this approach, the dataset is randomly partitioned into 10 equal-size subsets. The classifier is trained using 9 of the subsets, and the accuracy of its prediction is tested on the remaining subset. As part of the supervised learning, taxonomic labels are supplied for the DNA sequences in the 9 subsets used for training. The process is repeated 10 times. The accuracy score of the classifier is then computed as the average of the accuracies obtained in the 10 separate experiments.

### Classical Multidimensional Scaling (MDS)

Classical multidimensional scaling takes a pairwise distance matrix (*n × n* matrix, for *n* input items) as input and produces *n* points in a *q*-dimensional Euclidean space, where *q ≤ n −* 1. More specifically, the output is an *n × q* coordinate matrix, where each row corresponds to one of the *n* input items, and that row contains the *q* coordinates of the corresponding item-representing point [11]. The Euclidean distance between each pair of points is meant to approximate the distance between the corresponding two items in the original distance matrix.

These points can then be simultaneously visualized in a 2- or 3-dimensional space by taking the first 2, respectively 3, coordinates (out of *q*) of the coordinate matrix. The result is a Molecular Distance Map [43], and the MoDMap of a genomic dataset represents a visualization of the simultaneous interrelationships among all DNA sequences in the dataset.

### Software implementation

The algorithms for ML-DSP were implemented using the software package MATLAB R2017A, license no. 964054, as well as the open-source toolbox Fathom Toolbox for MATLAB [58] for distance computation. All software can be downloaded from https://github.com/grandhawa/MLDSP. The user can use this code to reproduce all results in this paper, and also has the option to input their own dataset and use it as training set for the purpose of classifying new genomic DNA sequences.

All experiments were performed on an ASUS ROG G752VS computer with 4 cores (8 threads) of a 2.7GHz Intel Core i7 6820HK processor and 64GB DD4 2400MHz SDRAM.

### Datasets

All datasets in this paper are at https://github.com/grandhawa/MLDSP under DataBase directory. The mitochondrial DNA dataset comprised all of the 7,396 complete reference mtDNA sequences available in the NCBI Reference Sequence Database RefSeq on June 17, 2017 (any letters “N” in these sequences were deleted). We performed computational experiments on several different subsets of this dataset. The dengue virus dataset contained all 4,721 dengue virus genomes available in the NCBI database on August 10, 2017.

For the performance comparison between ML-DSP and other alignment-free and alignment-based methods we also used the benchmark datasets of 38 influenza virus sequences, and 41 mammalian complete mtDNA sequences from [44].

## Author’s contributions

G.S.R. and L.K. conceived the study and wrote the manuscript. G.S.R. designed and tested the software. G.S.R., L.K. and K.A.H. conducted the data analysis and edited the manuscript, with K.A.H. contributing biological expertize. All authors read and approved the final manuscript.

### Acknowledgements

This work was supported by NSERC (Natural Science and Engineering Research Council of Canada) Discovery Grants R2824A01 to L.K., and R3511A12 to K.A.H. We thank Michael Pang for an independent Python implementation and reproducing some of the computational results, and Maximilian Soltysiak and Nicholas A. Boehler for comments on the manuscript and testing the software tool.

## Footnotes

1 For *m* clusters, the *m× m* confusion matrix has its rows labelled by the true classes and columns labelled by the predicted classes: The cell (*i, j*) shows the number of sequences that belong to the true class *i*, and have been predicted to be of class *j* (green indicates a correct prediction, and pink/red indicates incorrect predictions).

2 Note that if one changes the numerical representation from Integer to “Just-A”, the average accuracy of ML-DSP for the Mammalia dataset in Table 5 increases from 89.2% to 91.2%.

